# New Statistical Methods for Constructing Robust Differential Correlation Networks

**DOI:** 10.1101/393991

**Authors:** Danyang Yu, Zeyu Zhang, Kimberly Glass, Jessica Su, Dawn L. DeMeo, Kelan Tantisira, Scott T. Weiss, Weiliang Qiu

## Abstract

The interplay among microRNAs (miRNAs) plays an important role in the developments of complex human diseases. Co-expression networks can characterize the interactions among miRNAs. Differential correlation network is a powerful tool to investigate the differences of co-expression networks between cases and controls. To construct a differential correlation network, the Fisher’s Z-transformation test is usually used. However, the Fisher’s Z-transformation test requires the normality assumption, the violation of which would result in inflated Type I error rate. Several bootstrapping-based improvements for Fisher’s Z test have been proposed. However, these methods are too computationally intensive to be used to construct differential correlation networks for high-throughput genomic data. In this article, we proposed six novel robust equal-correlation tests that are computationally efficient. The systematic simulation studies and a real microRNA data analysis showed that one of the six proposed tests (ST5) overall performed better than other methods.

## Introduction

A microRNA (miRNA) is a non-coding RNA molecule that plays an important role in RNA silencing and post-transcriptional regulation of gene expression.^1,2^ There is increasing evidence that miRNAs are closely related to various human complex diseases. Each miRNA can interact with hundreds of genes and plays various roles in tumorigenesis, metastasis, proliferation.^3^ In addition, a single gene can also be targeted by multiple miRNAs, which constitutes complex miRNA-target interactions. ^3^

The interplay among miRNAs plays an important role in the development of complex human diseases. The network provides a natural way to model the interactions among miRNAs, with nodes representing miRNAs and edges representing interactions between miRNAs.^4^ The main advantages of network-based approaches include their feasibility for large amounts of data, robustness to disturbances, and ease of visual interpretation.^5^ The development of network theory makes it possible to calculate the global properties of these networks, providing insight into the behavior of the systems they represent.^6^ A gene co-expression network is an undirected graph in which nodes represent genes or probes, and a pair of nodes is connected by an edge if there is a significant co-expression relationship between them.^7^ An edge weight indicates the magnitude of the co-expression (e.g., correlation) between the pair of nodes connected by the edge. A gene co-expression network focuses on the interplay of multiple genes, checking whether these genes are over or under expressed simultaneously.

By comparing the gene network based on diseased samples (cases) and that based on non-diseased samples (controls), one can know which gene pairs are involved in the development of the disease. We call a network of genes as a gene differential correlation network if the edges in the network connect pairs of genes having significantly different edge weights (i.e., correlations) between the gene co-expression network based on cases and that based on controls (Figure 1). To the best of our knowledge, no research has constructed differential correlation networks using miRNAs yet.

**Figure 1.**
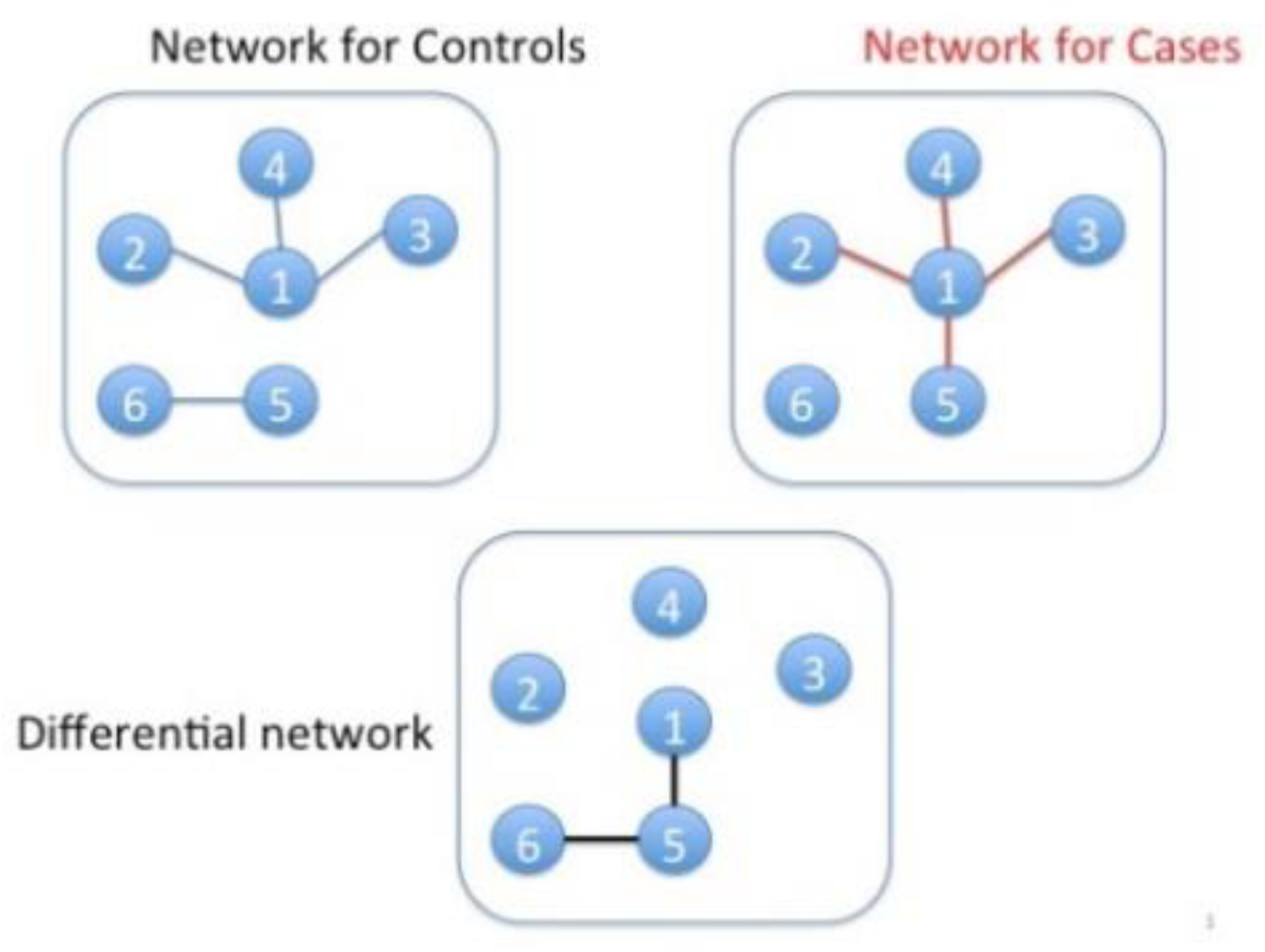
An illustration of a differential correlation network.

To construct a differential correlation network, we usually first test for each pair of nodes if the correlation between the pair is the same or not between cases and controls. We then connect a pair of nodes if the difference of the correlations between cases and controls is greater than a threshold. The benchmark statistical method for testing equal correlation of a pair of random variables between two independent populations is the Fisher’s Z-transformation test. However, it is sensitive to the violation of the normality assumption. The normal distribution cannot be guaranteed in real data analysis. The violation of the normality assumption would result in inflating type I error rate (i.e., false positive rate).

Some improved tests for equal correlation have been proposed to be robust against the violation of the normality assumption. However, there are some limitations of those methods. For instance, the two methods (twocor and twopcor) proposed by Wilcox (2011)^8^ are bootstrapping based methods, which are computationally intensive. To construct a gene differential correlation network, we need to test equal correlation between cases and controls for G(G-1)/2 pairs of gene probes, where G is the number of gene probes, which is usually large (∼20,000) in whole genome-wide data analysis. Hence, it is not efficient to use bootstrapping-based methods to construct gene differential correlation networks.

ROS-DET (combination of robust correlations and hypothetical testing) proposed by Kayano et al. (2011) only focuses on pairs of genes that have positive correlations in one subject group and negative correlations in another subject group.^9^ It ignores the scenarios where two correlations are significantly different but have the same directions. The result of other methods without bootstrapping, such as Zou’s method^10^ and HC4 (heteroscedastic-consistent estimators) method^11^, can be unsatisfactory, even under normality assumption.^12^ Hence, there is a great need to develop a robust and fast test for equal correlation. In this article, we proposed 6 novel tests for equal correlation and performed systematic simulation studies and a real data analysis to compare the performances of these new methods with existing methods.

## Results

### Simulation studies

We compared the 6 proposed equal-correlation tests (ST1, ST2, ST3, ST4, ST5, ST6) with 4 existing tests (twopcor, twocor, twohc4cor^8^and Fisher’s Z-transformation test) using systematic simulation studies, in which we evaluated if the 6 proposed methods could achieve higher power than the 4 existing methods, while keeping the nominal type I error rate (0.05), when random variables *X* and *Z* are generated from normal or non-normal distributions. The methods twopcor and twocor are bootstrapping-based methods.

Following Wilcox (2009), we generated the observations of the random variables *X* and *Z* from g-and-h-distributions^13^ (see Section 3 of Supplementary Document I for the definition of a g-and-h distribution) for cases and controls. In a g-and-h distribution, the parameters g and h are both non-negative. If both g and h are equal to zero, then the g-and-h distribution is the standard normal distribution. A positive value of g indicates a skewed distribution. A positive value of h indicates that the g-and-h distribution has heavier tail than the standard normal distribution. As g becomes larger, the g- and-h distribution would be more asymmetric. As h becomes larger, the g- and-h distribution would have heavier tail than the standard normal distribution. We considered four scenarios: (1) g=0.2 and h=0.2 (asymmetrical distribution with heavy tail); (2) g=0.2 and h=0 (asymmetrical distribution with relatively light tail); (3) g=0 and h=0.2 (symmetrical distribution with heavy tail); and (4) g=0 and h=0 (standard normal distribution).

To generate observations for a pair of random variables *X* ad *Z*, we first generated random numbers *X1* for cases and *X2* for controls. Then we generated random numbers of *Z1* and *Z2* via the formula z_1_= *θ*_1_*x*_1_ +*λ_j_*(x_1_)*e*_1_for cases and z_2_ = *θ*_2_*x*_2_ + *λ_j_*(*x*_2_)*e*_2_for controls. The choices for *λ_j_*(*x*_*i*_), *i =*1, 2 were taken to be 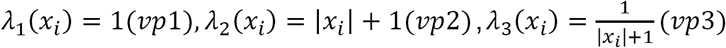 respectively, which indicate different patterns of the variance of *Z*. The random error terms *e*_1_ and *e*_2_ have same distributions as *x*_1_ and *x*_2_.^12^

To evaluate the effect of sample size on the performances of the equal-correlation tests, we considered 3 different sample sizes: (1) 30 cases and 30 controls; (2) 100 cases and 100 controls; and (3) 200 cases and 200 controls. To evaluate the type I error rates of the tests, we considered two scenarios: (1) *θ*_1_ = *θ*_2_ = 0; and (2) *θ*_1_ = *θ*_2_ = 1; To evaluate the powers of the tests, we set *θ*_1_ = 0 and *θ*_2_ = 1. So we totally have 4 × 3 × 3 × 2 = 72 scenarios for evaluating Type I error rates and 4 × 3 × 3 × 1 = 36 scenarios for evaluating powers.

For each scenario, we generated 100 datasets. For each dataset, we generated 1000 pairs of random variables X and Z for cases and controls, respectively. Because bootstrapping-based methods (twopcor and twocor) would cost too much time, we evaluated the performances of twopcor and twocor only in the scenarios where nCases=nControls=100, g=0.2, and h=0.2 (nCases is the number of cases and nControls is the number of controls).

To evaluate the performances of equal-correlation tests, we used Type I error rate and power in simulation studies. Figure 3 and Figure S4 showed that the median of Type I error rates of twocor and ST5 did not exceed the nominal level 0.05 in all scenarios in the simulation studies, which shows the excellent robustness of twocor and ST5. The medians Type I error rates of ST1 and ST2 (when h=0, *θ*_1_ = 0, *θ*_2_ = 0), and ST6 (when variation pattern is vp3, h=0,*θ*_1_ = 1, *θ*_2_ = 1 and n=100 or 200) were just a little bit higher than 0.05 in few scenarios, which shows the symmetry does not affect the type I error rates of ST1, ST2 and ST6, but the heave tail does. For twopcor, the medians of Type I error rates were higher than 0.05 when the variance pattern was vp1 or vp2. When *θ*_1_ = 1, *θ*_2_ = 1 (i.e., correlations between X and Z are non-zero, but the same, in both cases and controls), the median of Type I error rates of twohc4cor and ST4 are higher than 0.05; For ST3 the median of Type I error rates are higher than 0.05 when *θ*_1_ = 0, *θ*_2_ = 0 (i.e., correlations between X and Z are zero in both cases and controls).

**Figure 2:**
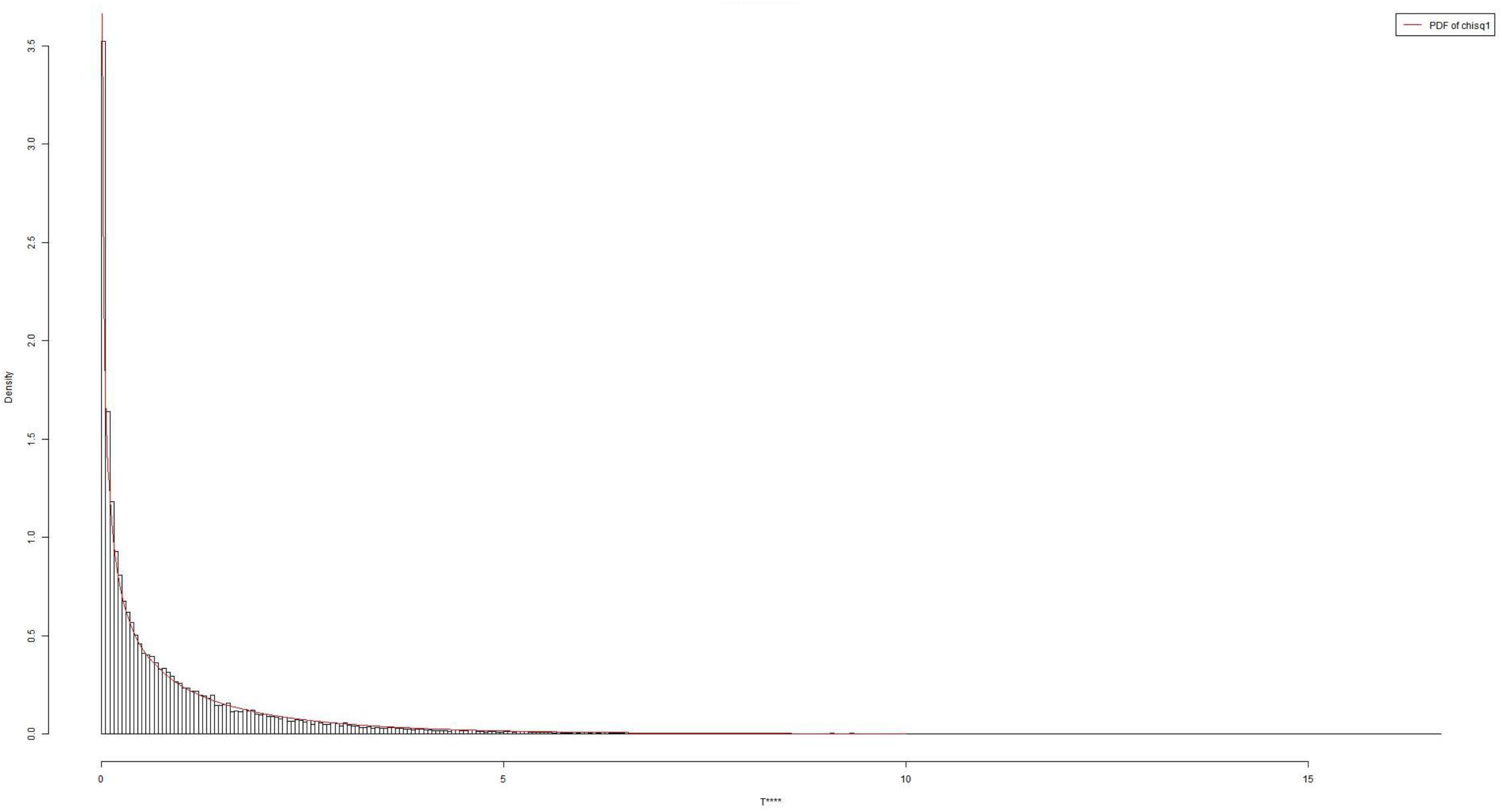
histogram of *T*^*V*^.

**Figure 3:**
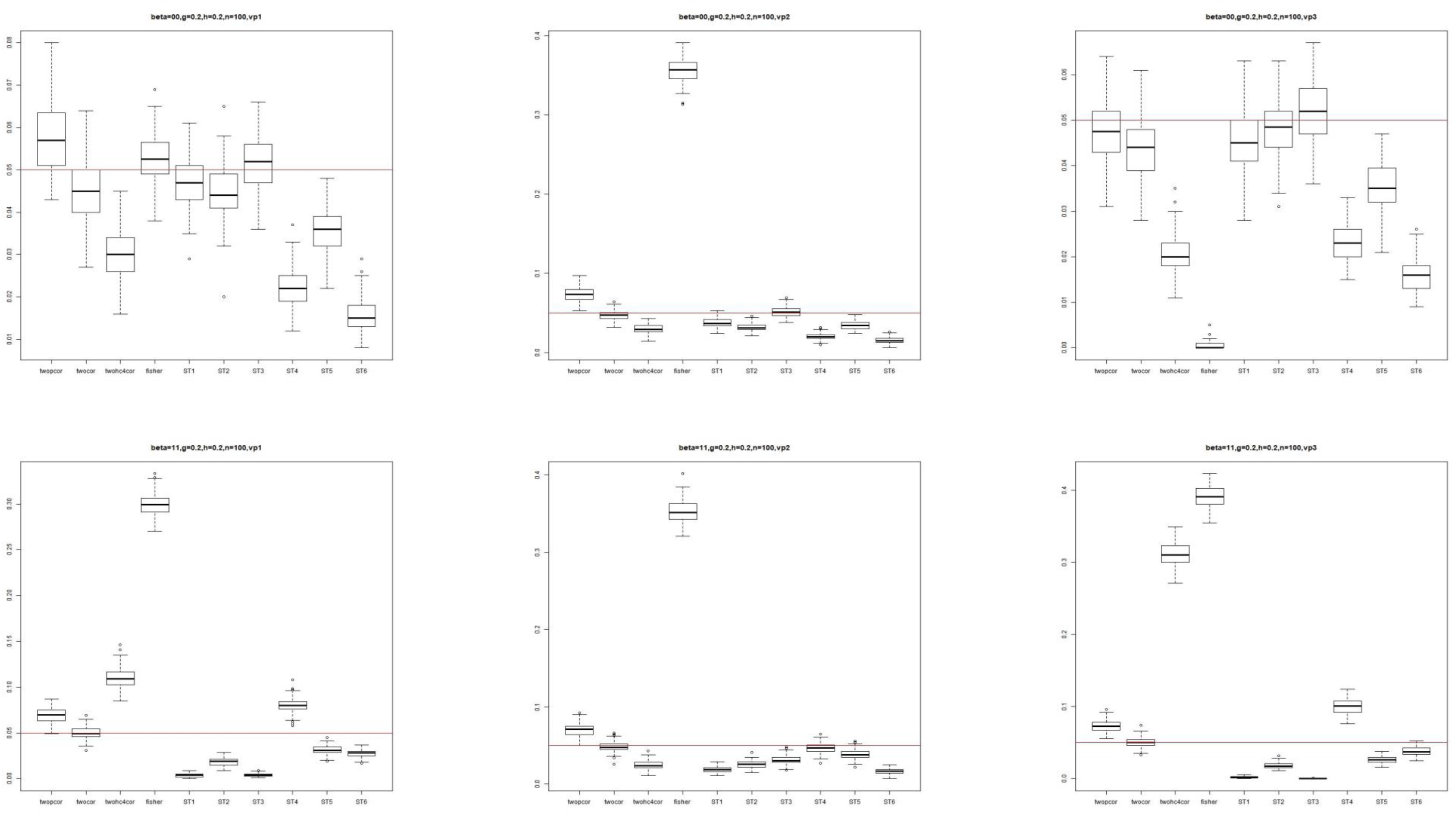
boxplots of type I error rates.

For the power analyses, the median powers of all six methods increase as sample size increases. For all six methods, the powers were smaller when the variation pattern was vp2. Among the methods (ST1, ST2, ST5, ST6 and twocor) that have the median Type I error rates smaller than 0.05 in almost all simulation scenarios, the ST5 and twocor always have the highest power, which is not affected by sample sizes, distributions and variance patterns. The median powers of ST5 were almost equal to those of twocor in all scenarios.

We summarized the simulation results in Figure 5 and Table S1. Figure 5 is a plot of the number *n*_*reject*_of scenarios with mean type I error rate significantly >0.05 versus the median rank of power *m*. For each scenario and each equal-variance test, we used one-sample t-test to test the null hypothesis *H*_0_ that the mean of the 100 estimated type I error rates from 100 simulated data set is significantly ≤ 0.05 versus the alternative hypothesis *H*_a_ that the mean of the 100 estimated type I error rates from 100 simulated data set is significantly >0.05. If the p-value of the one-sample t-test < 0.05, we claimed for this scenario and this equal-variance test, the mean type I error rate > 0.05. For a given scenario, we ranked in terms of power the equal-variance tests that did not reject the null hypothesis *H*_0_. For ranks with ties, average ranks were used. For the equal-variance tests that rejected the null hypothesis *H*_0_, we set their ranks as missing values. Since for each scenario evaluating power (*θ*_1_ = 0, *θ*_2_ = 1), there are two corresponding scenarios evaluating Type I error rate (*θ*_1_ = *θ*_2_ = 0 or *θ*_1_ = *θ*_2_ = 1), we set ranks as missing values if one of the two corresponding scenarios with mean Type I error rate > 0.05. We then obtained the median *m* of the rank for each equal-variance test. The left panel of Figure 5 is based on all scenarios. The proposed equal-variance test ST5 located at the left-bottom corner of the left panel of Figure 5 (*n*_*reject*_ = 0, *m =*1.75), indicating it had the smallest number of false positive rate and largest power, hence performed best. The methods twocor and twopcor have not appeared in the left panel of Figure 5 because they are bootstrapping-based methods, which are computationally intensive. We only include them in the scenarios where nCases=nControls=100, g=0.2, and h=0.2. The right panel of Figure 5 is the plot for these scenarios, from which we can see that twocor performed the best (*n*_*reject*_ = 0, *m =* 1.5) and ST5 performed the second (*n*_*reject*_ = 0, *m =* 2).

**Figure 4:**
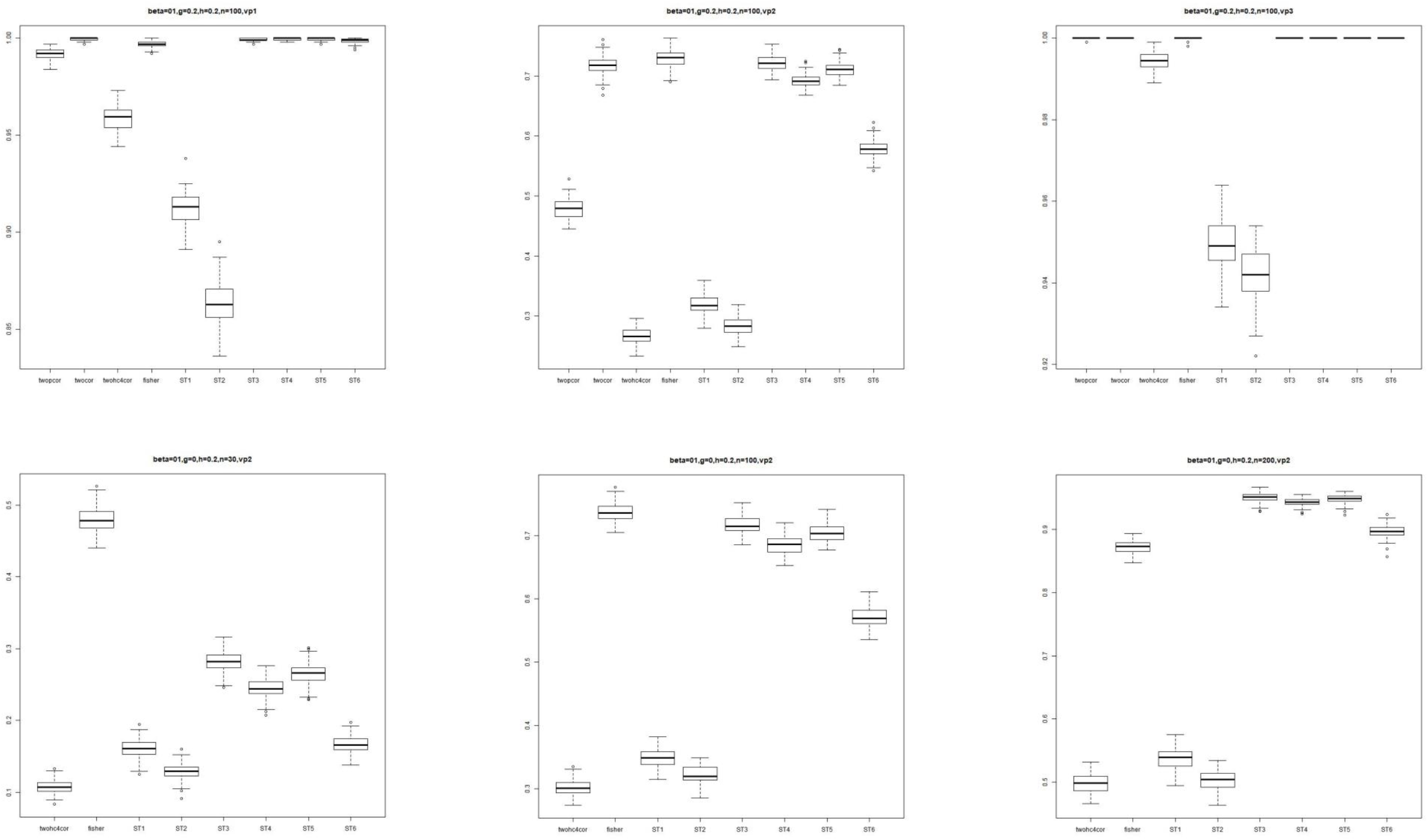
boxplots of powers.

**Figure 5:**
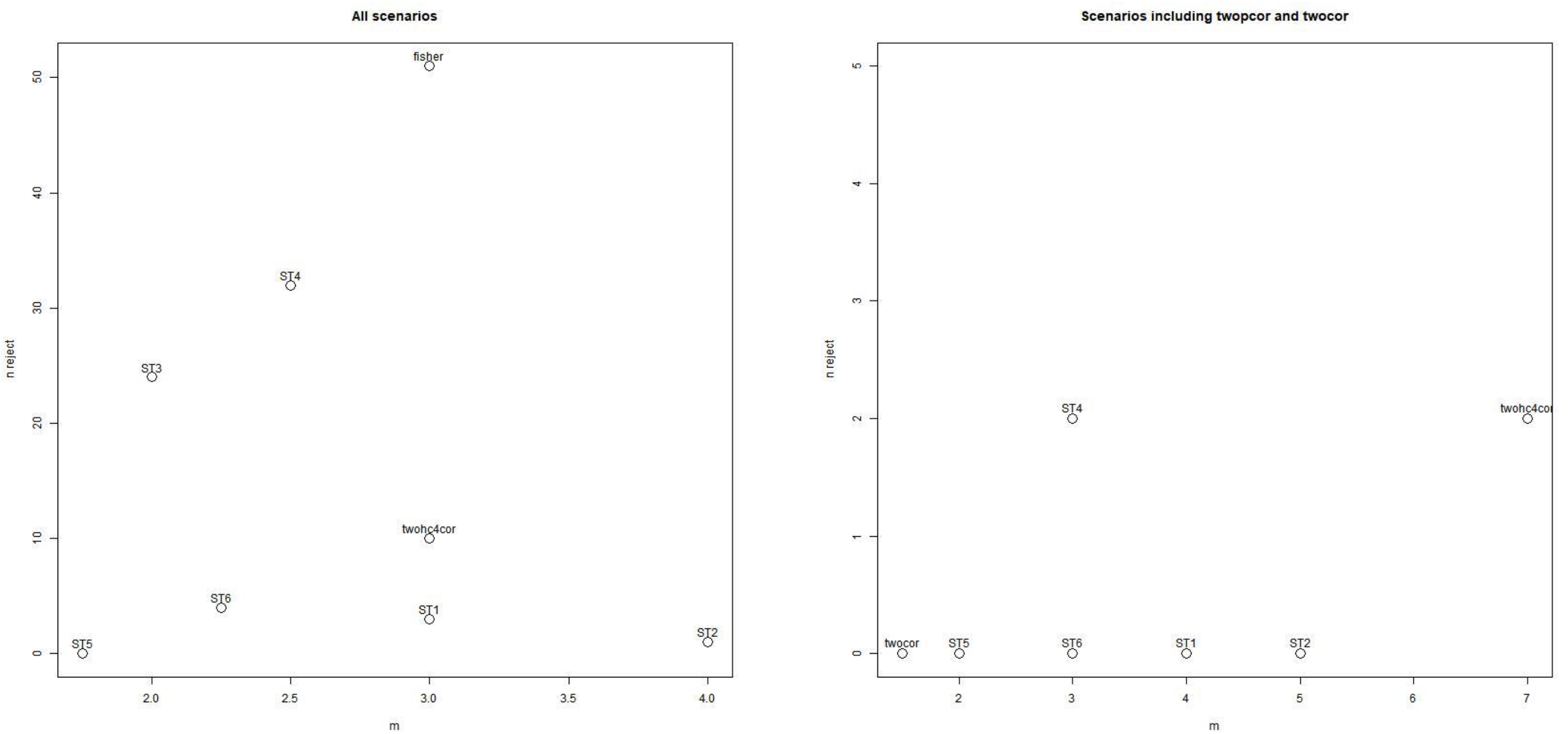
*n*_*reject*_ is the number of scenarios with type I error rate > 0.05, m is the median ranks of power. We set ranks missing value if one of the two corresponding type I error rates (two scenarios with same sample size, variation pattern, g and h) of the power is larger than 0.05. The left panel was obtained based on all scenarios, the right panel was obtained based on scenarios including twopcor and twocor.

### Real data analysis

We downloaded the miRNA dataset GSE15008 (https://www.ncbi.nlm.nih.gov/geo/query/acc.cgi?acc=gse15008) from the public data repository Gene Expression Omnibus (GEO, https://www.ncbi.nlm.nih.gov/geo) to construct differential correlation networks of miRNAs using the 6 equal-correlation tests (3 proposed tests and 3 existing tests). GSE15008 contains 677 miRNAs from 174 non-small-cell lung cancer (NSCLC) tissues and 187 adjacent normal tissues from patients. After data preprocessing, 178 miRNAs were kept for differential correlation analysis (The details about the data preprocessing are shown in Section 4 of Supplementary Documents I). The quantile plots (Figure S2) and plots of the top 2 principal components (Figure S3) based on the 178 miRNAs showed no obvious patterns.

We randomly divided the 174 NSCLC samples into two equal parts. One part (87 samples) formed the cases of the discovery set and the other part (87 samples) formed the cases of the validation set. Similarly, we randomly divided the 187 normal samples into roughly two equal parts. One part (94 samples) formed the controls of the discovery set and the other part (93 samples) formed the controls of the validation set.

We compared ST1, ST5, ST6, Fisher’s test, twohc4cor, and twocor in the real data analysis. We did not include ST2 because of its low power in the simulation studies. We did not include ST3 and ST4 because they were no better than ST5 and ST6 in the simulation studies. We did not include twopcor because of its long computational time and its low power in the simulation studies.

We claimed that the differential correlation of a pair of miRNAs is validated if its FDR-adjusted p-value < 0.05 in the discovery set and raw p-value < 0.05 in the validation set. We applied Benjamini and Yekutieli method^14^ to calculate FDR-adjusted p-values.

We called the network formed by the pairs of miRNAs with validated differential correlations as the validated differential correlation network. We visualize validated differential correlation networks by Cytoscape.

We chose the miRNA having the maximum number of edges in the validated differential correlation network as the hub miRNA. We applied the web-tool miRSystem^15^ to predict the genes targeted by the hub miRNA and to obtain the KEGG pathways enriched in these genes.

To evaluate the differential correlation networks in the real data analysis, we used the proportion of the validated edges in the discovery set [in eq], where *e*_1_ is the number of validated edges and *e*_2_ is the number of edges detected based on the discovery set.

The numbers *e*_1_ of validated edges obtained by the 6 proposed tests are 0 (ST1), 35 (ST5), 37 (ST6), 71 (Fisher), 0 (twohc4cor) and 440 (twocor). ST6 had the highest validation rate (r=100.00%), followed by ST5 (r=94.59%), Fisher (r=88.75%), and twocor (r=74.83%). In terms of running time, *Fisher* is the fastest method among the 6 methods, which took only 2.42 seconds. ST1 is the second fastest method, which took 5.97 seconds. ST5 and ST6 took 35.92 seconds and 37.32 seconds, respectively. Twohc4cor used 89.88 seconds, which is around 2.5 more times than ST5/ST6. Twocor, the bootstrapping-based method, took the longest time: 31429.21 seconds (i.e., 8.73 hours). The results of the differential correlation analyses of the 6 methods for the real dataset GSE15008 are summarized in Table 1.

**Table 1.** Summary of the differential correlation analyses of the 6 methods for the real dataset GSE15008

|  | ST1 | ST5 | ST6 | Fisher | twohc4cor | twocor |
| --- | --- | --- | --- | --- | --- | --- |
| $e_1$ | 0 | 35 | 37 | 71 | 0 | 395 |
| $e_2$ | 0 | 37 | 37 | 80 | 0 | 537 |
| $\frac{e_1}{e_2}$ | / | 94.59% | 100.00% | 88.75% | / | 73.56% |
| Hubs | / | hsa-miR-143 | hsa-miR-143 | hsa-miR-492 | / | hsa-miR-363 |
| Running time(seconds) | 5.97 | 35.92 | 37.32 | 2.42 | 89.88 | 31429.21 |

Figure 6 showed the validated differential correlation networks based on ST5, ST6, Fisher, and twocor. The hubs in the differential correlation networks obtained by ST5 and ST6 are the same, i.e., hsa-miR-143, which targets 531 genes (Table S2). These 531 genes are enriched in 5 KEGG pathways (LEISHMANIASIS, PRION DISEASES, GNRH SIGNALING PATHWAY, ECM RECEPTOR INTERACTION, and ALZHEIMER’S DISEASE) (Table S3). The hub detected by *Fisher* is has-miR-492, which targets 107 genes (Table S2). These 107 genes are not enriched in KEGG pathways. The hub detected by twocor is hsa-miR-363, which targets 573 genes. These 573 genes are enriched in 5 KEGG pathways (DILATED CARDIOMYOPATHY, CALCIUM SIGNALING PATHWAY, ECM RECEPTOR INTERACTION, LONG TERM POTENTIATION, and REGULATION OF ACTIN CYTOSKELETON) (Table S3).

**Figure 6:**
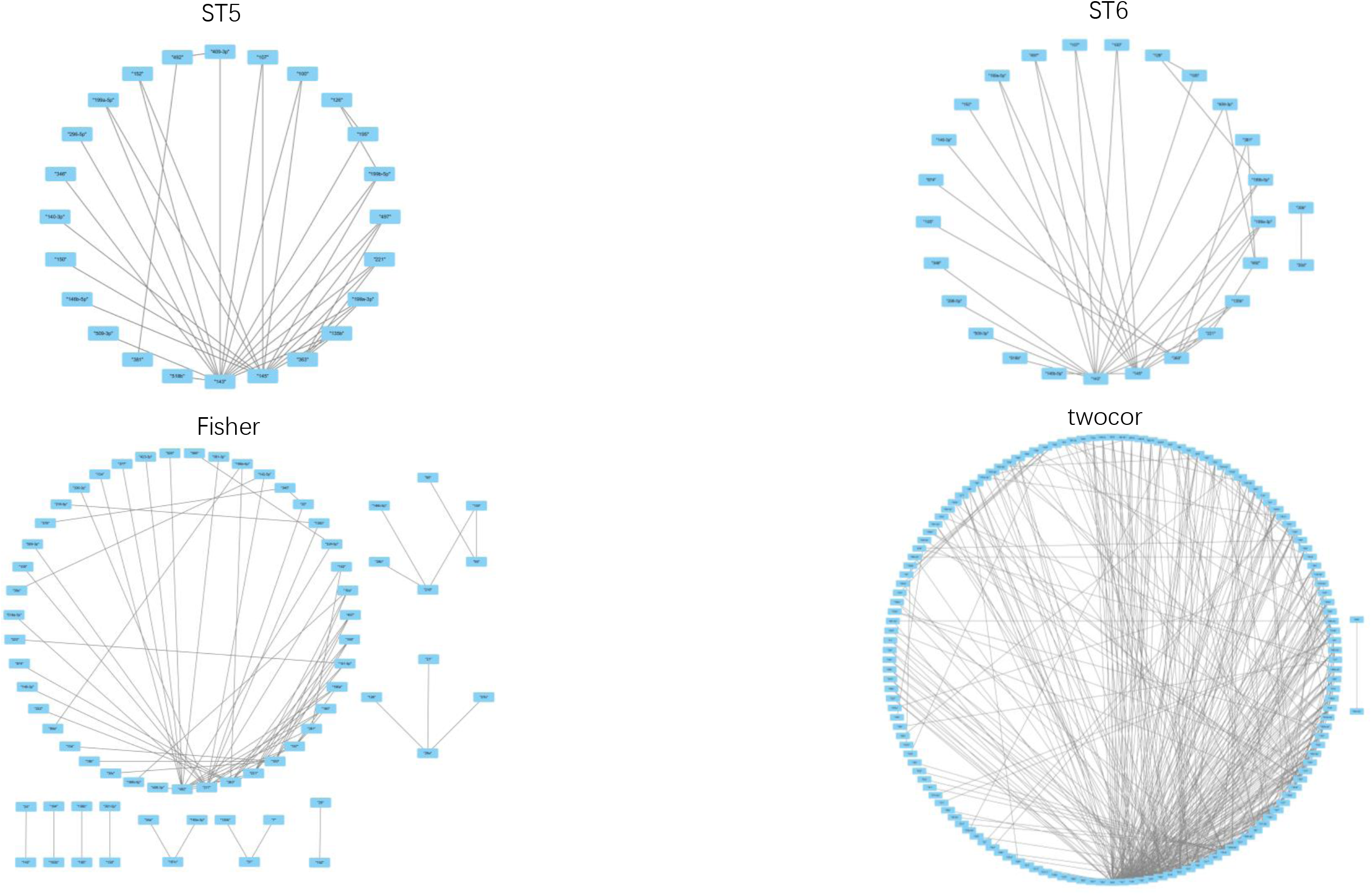
validated differential correlation network.

## Discussion

A differential correlation network of miRNAs, characterizing the differences between the co-expression network among cases and that among controls, could help understand how pairs of miRNAs affect the disease of interest. In this article, we proposed six novel robust tests (ST1, ST2, ST3, ST4, ST5, and ST6) for equal correlation and compared them with four existing tests (twocor, twopcor, Fisher’s Z-transformation test, and twohc4cor) in the construction of differential correlation networks. Simulation studies showed that ST5 performed as well as twocor (a bootstrapping-based method), and had the highest powers among the tests that kept the nominal type I error rates in all scenarios in our simulation studies. Real data analysis also showed the good performance of ST5. ST5 had the second highest validation rate (r=94.59%) in the real data analysis, followed by Fisher (r=88.75%) and twocor (r=74.83%), while ST6 had the highest validation rate (r=100%). Furthermore, ST5 is computationally fast. Hence, the proposed equal-correlation test ST5 could be used to construct robust differential correlation networks in genomic data analysis.

Although twocor showed good performance in simulation studies, twocor detected too many differential correlation edges (*e*_1_ = 395, *e*_2_=537), but had the smallest validation rate (r=73.56%) in the real data analysis. Moreover, as a bootstrapping-based method, twocor is computationally intensive. Therefore, twocor probably is not suitable to construct differential correlation networks in genomic data analysis, in which the number of nodes is large.

In the real data analysis, the rank of the proportion of validated edges is ST6, ST5, Fisher, and twocor. ST1 and twohc4cor failed to detect any significant differential correlation based on the discovery sets. It indicates that ST1 and twohc4cor are not powerful methods, which are also shown in the simulation studies. In the validated differential correlation networks of ST5, ST6 and twocor, the top three nodes having the largest number of edges are same (hsa-miR-143, hsa-miR-145, hsa-miR-363). The miRSystem predicted that hsa-miR-143 (the hub detected by ST5 and ST6), has-miR-145, and hsa-miR-363 (the hub detected by twocor) are connected to pathways related to lung cancer, such as GNRH signaling pathway^16^ (empirical-p-value is 0.01490), calcium signaling pathway^17^ (empirical-p-value is 0.01872), and TGF-BETA signaling pathway^18^ (empirical-p-value is 0.02706). However, hsa-miR-492 (detected by Fisher) is not related to lung cancer (Table S3). We surmised the reason why the hub selected by Fisher is not related to lung cancer is the high false positive rates of the Fisher’s Z-transformation test.

In this article, there are a couple of limitations. Firstly, the sample size (174 NSCLC tissues and 187 normal tissues) of the GEO dataset GSE15008 is not very large and we did not find an independent dataset to do validation. Instead, we randomly split the GSE15008 dataset into two sets: discovery set and validation set. Secondly, we could not derive the asymptotic or approximate distribution of the ST5 test statistic yet. Instead, we numerically demonstrated that the distribution of the ST5 test under the null hypothesis could be approximated by the chi square distribution with one degree of freedom. Further research on the asymptotic distribution of ST5 is needed.

In summary, we proposed 6 robust tests for equal correlation to construct differential correlation networks and found ST5 had overall good performance in both the simulation studies and the real data analysis. The ST5 test can be used to construct differential correlation network to characterize the association between the interactions between genes (not limited to mi-RNAs) to diseases (not limited to lung cancer), hence to help uncover the molecular mechanisms of complex human diseases.

## Methods

To construct a differential correlation network for a set of miRNAs, we first test for each pair of miRNAs if their correlation among cases is the same as that among controls using an equal-correlation test. We then set a criterion to determine if the test is significant or not (i.e., if the pair of miRNAs should be connected by an edge in the differential correlation network or not). To determine if a test is significant or not, we need to control for multiple testing since G(G-1)/2 tests are performed, where G is the total number of miRNAs. We claimed that the differential correlation of a pair of miRNAs is validated if its FDR-adjusted p-value < 0.05 in the discovery set and raw p-value < 0.05 in the validation set.

### Novel equal-correlation tests

For given two random variables X and Z, we would like to test if the correlation corr(X, Z) in cases is the same as that in controls. Inspired by the joint test of equal mean and equal variance proposed by Ahn and Wang^19^ and the improved Ahn and Wang’s equal-variance tests proposed by Qiu et al.^20^, we proposed 6 robust tests (ST1, ST2, ST3, ST4, ST5, ST6) for equal correlation without the normality assumption.

### ST1 Test

Let’s consider the following logistic regression:

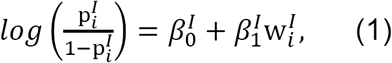

were

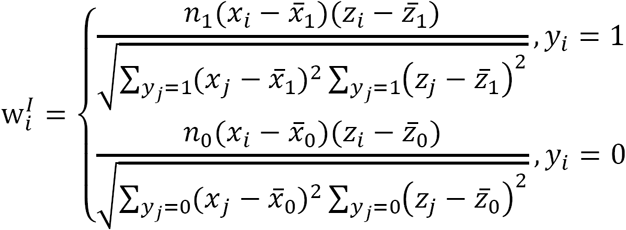

*n*_0_ represents the number of controls, *n*_1_ represents the number of cases, *y_i_ =* 1 indicates that the *i*-th subject is a case, and *y_i_ =* 0 indicates that the *i*-th subject is a control.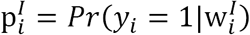is the probability that the *i*-th subject is a case given 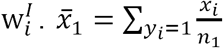and 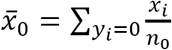are sample means of *X* in cases and controls, respectively.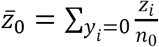and 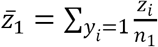are sample means of *Z* in cases and controls, respectively. The score test statistic of the above logistic regression for testing the null hypothesis 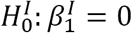is

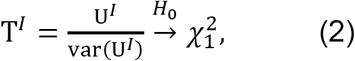

where 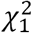 is the chi squared distribution with one degree of freedom,

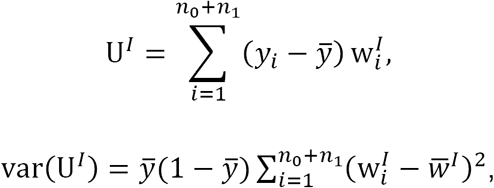

And

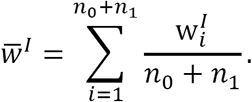

That is, when U^*I*^ is large, we reject 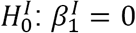. We can show that 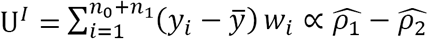(See the section 1 of Supplementary Document I).

Hence, testing 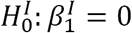is equivalent to test for *ρ*_1_ = *ρ*_2_.

Note that in logistic regression (1), the random variable *y*_*i*_ is conditional on the random variable 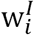.

We denote the score test in (2) as ST1 Test.

### ST2 Test

ST2 test is an improved version of ST1 by replacing the sample variances by the square of median absolute deviations (MAD) since MAD is more robust to outliers than standard deviation. Let’s consider the following logistic regression:

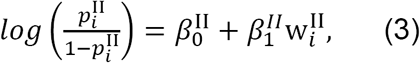

where 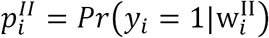

The related formulas are shown below:

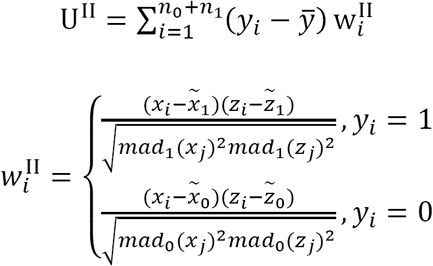

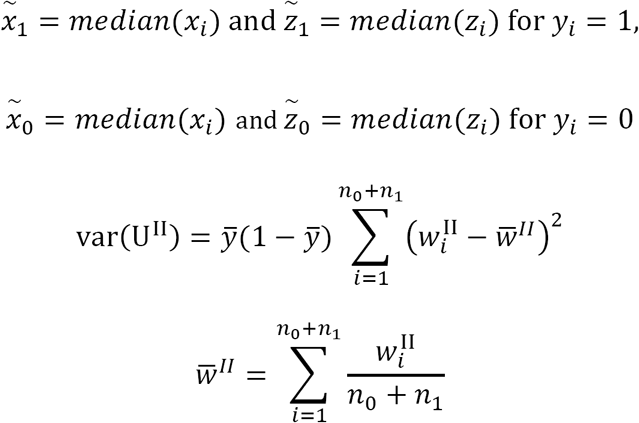

The ST2 test statistic for testing the null hypothesis 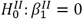is

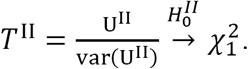

### ST3 Test

To get more robust weights than 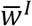and 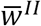, we utilized the M type correlation^8^ in ST3 test. The idea of the M type correlation is to use a robust version of trimmed mean to replace sample mean in calculating sample correlation. Please refer to Section 2 of Supplementary Document I for the details about the ST3 test.

### ST4 Test

In ST4 test, we defined the weights 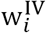based on the Spearman’s rank correlation. The formulas are below:

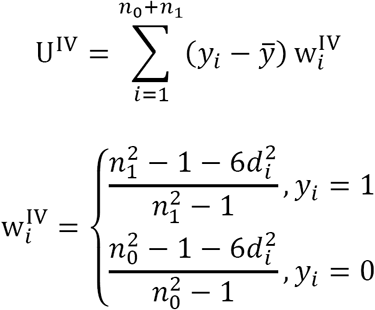

where *d_i_ =* rank(*x*_*i*_) - rank(z_*i*_), where rank (X) represents the ranks of x in both case and control, and

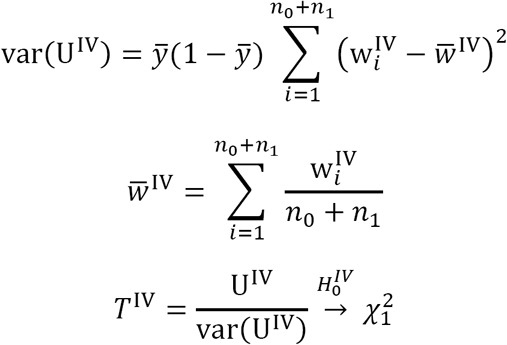

### ST5 Test

Our systematic simulations (see the Result Section) showed that the performances of ST3 and ST4 are complement of each other. So, we combined the test statistics of ST3 and ST4 to form the test statistic of ST5. Specifically, the test statistic of ST5 is the average of *T*^III^ and *T*^IV^:

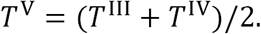

It is challenging to derive the asymptotic distribution of the test statistic *T*^*V*^. We guess that the asymptotic distribution of the test statistic *T*^*V*^ is close to the chi square distribution with one degree of freedom 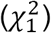. To numerically support this guess, we generated a simulated dataset with 50,000 pairs of random variables X and Z for 100 cases and 100 controls from a g-and-h distribution under the null hypothesis 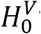that the correlation between *X* and *Z* in cases is the same as that in controls (see Simulations Section for more details). We then calculated the values of *T*^*V*^ for each of the 50,000 pairs of random variables *X* and *Z* and drew the histogram of these 50,000 values of *T*^*V*^. Figure *2* showed that the histogram of the test statistic *T*^*V*^ is very close to the density of the chi-squared distribution with one degree of freedom. Hence, in this article, we assumed that 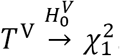 Further investigation is warranted.

### ST6 Test

We can combine ST3 and ST4 based on the following multiple logistic regression:

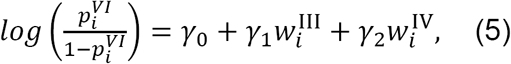

where 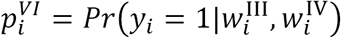The score test statistic for testing for the composite null hypothesis 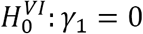and γ_2_ = 0 is

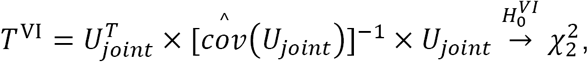

Where

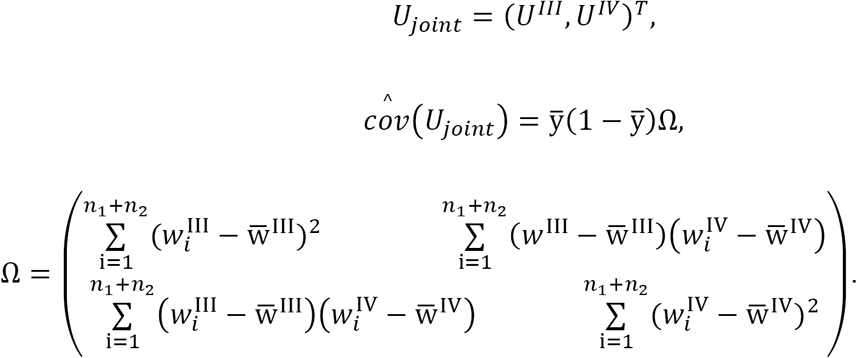

The proof that the asymptotic distribution of 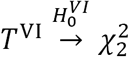is given in the Supplementary Document II.

## Author contributions

W.Q. initiated and supervised the project. D.Y., assisted by W.Q., designed and implemented the simulation study and real data analysis. Z.Z., K.G., J.S., D.D., K.T., and S.W. provided comments and suggestions to the project, helped interpret the results, and design the figures and tables. All authors contributed to the manuscript writing and approved the submission of the manuscript.

## Materials & Correspondence

The contact information of the corresponding author is as follows: Dr. Weiliang Qiu, Channing Division of Network Medicine, Brigham and Women’s Hospital/Harvard Medical School, 181 Longwood Avenue, Boston, MA 02115 USA. Tel: 1-617-525-0841.

## Data Availability

The datasets analyzed during the current study are available in the GEO repository, https://www.ncbi.nlm.nih.gov/geo/query/acc.cgi?acc=gse15008.

## Code availability

The R package including functions for implementing ST1, ST2, ST3, ST4, ST5, ST6, and Fisher’s Z-transformation test can be download from the website: https://sites.google.com/a/channing.harvard.edu/weiliang-qiu/Home/software or https://CRAN.R-project.org/package=corTest.

## References

1. Bartel, D. P. MicroRNAs: Genomics, Biogenesis, Mechanism, and Function. Cell 116, 281–297 (2004).

2. Ambros, V. The functions of animal microRNAs. Nature (2004). doi:10.1038/nature0287.

3. Chen, X., Xie, D., Zhao, Q. & You, Z.-H. MicroRNAs and complex diseases: from experimental results to computational models. Brief Bioinform doi:10.1093/bib/bbx13.

4. Barabási, A.-L. & Oltvai, Z. N. Network biology: understanding the cell’s functional organization. Nature Reviews Genetics 5, 101–113 (2004).

5. Silverman, E. K. & Loscalzo, J. Network medicine approaches to the genetics of complex diseases. Discov Med 14, 143–152 (2012).

6. Glass, K. & Girvan, M. Finding New Order in Biological Functions from the Network Structure of Gene Annotations. PLOS Computational Biology 11, e1004565 (2015).

7. Stuart, J. M., Segal, E., Koller, D. & Kim, S. K. A Gene-Coexpression Network for Global Discovery of Conserved Genetic Modules. Science 302, 249–255 (2003).

8. Wilcox, R. R. Introduction to Robust Estimation and Hypothesis Testing. (Academic Press, 2011).

9. Kayano, M., Takigawa, I., Shiga, M., Tsuda, K. & Mamitsuka, H. ROS-DET: robust detector of switching mechanisms in gene expression. Nucleic Acids Res 39, e74 (2011).

10. Zou, G. Y. Toward using confidence intervals to compare correlations. Psychol Methods 12, 399–413 (2007).

11. Cribari-Neto, F. Asymptotic inference under heteroskedasticity of unknown form. Computational Statistics & Data Analysis 45, 215–233 (2004).

12. Wilcox, R. Comparing Pearson Correlations: Dealing with Heteroscedasticity and Nonnormality. Communications in Statistics - Simulation and Computation 38, 2220–2234 (2009).

13. Hoaglin, D. C. Summarizing Shape Numerically: The g-and-h Distributions. in Exploring Data Tables, Trends, and Shapes (eds. Hoaglin, D. C., Mosteller, F. & Tukey, J. W.) 461–513 (John Wiley & Sons, Inc., 2006). doi:10.1002/9781118150702.ch1.

14. Benjamini, Y. & Yekutieli, D. The control of the false discovery rate in multiple testing under dependency. Ann. Statist. 29, 1165–1188 (2001).

15. Lu, T.-P. et al. miRSystem: An Integrated System for Characterizing Enriched Functions and Pathways of MicroRNA Targets. PLOS ONE 7, e42390 (2012).

16. Lu, C., Huang, T., Chen, W. & Lu, H. GnRH participates in the self-renewal of A549- derived lung cancer stem-like cells through upregulation of the JNK signaling pathway. Oncology Reports 34, 244–250 (2015).

17. Yang, H., Zhang, Q., He, J. & Lu, W. Regulation of calcium signaling in lung cancer. J Thorac Dis 2, 52–56 (2010).

18. Jeon, H.-S. & Jen, J. TGF-beta signaling and the role of inhibitory Smads in non-small cell lung cancer. J Thorac Oncol 5, 417–419 (2010).

19. Ahn, S. & Wang, T. A powerful statistical method for identifying differentially methylated markers in complex diseases. in Biocomputing 2013 69–79 (WORLD SCIENTIFIC, 2012). doi:10.1142/9789814447973_000.

20. Qiu, W. et al. New Score Tests for Equality of Variances in the Application of DNA Methylation Data Analysis. 8 (2017).

